# PIEZO1 force sensing controls global lipid homeostasis

**DOI:** 10.1101/2023.07.23.550198

**Authors:** Laeticia Lichtenstein, Chew W Cheng, Elizabeth L Evans, Hannah J Gaunt, Fiona Bartoli, Eulashini Chuntharpursat-Bon, Shaili Patel, Charalampos Konstantinou, T Simon Futers, Melanie Reay, Gregory Parsonage, Justine Bertrand-Michel, Piruthivi Sukumar, Lee D Roberts, David J Beech

**Affiliations:** School of Medicine, University of Leeds, Leeds, LS2 9JT, UK; Department of Hepatobiliary and Transplant Surgery, St James’s University Hospital, Leeds, LS9 7TF, UK; MetaToul-Lipidomics Facility, INSERM UMR1048, Toulouse, France; Institut des Maladies Métaboliques et Cardiovasculaires, UMR 1297/I2MC, INSERM, Toulouse, France

## Abstract

How cardiovascular activity beneficially regulates lipid homeostasis is unclear. Here we hypothesise a mechanism in which mechanical force sensed by PIEZO1 ion channels in endothelium links blood flow to lipid regulation. We engineered mice for conditional deletion of PIEZO1 in endothelium and determined consequences for lipid regulation. Prominent are upregulated expression of hepatic *Cyp7a1* and intestinal *Ldlr* genes, which are pivotal in cholesterol catabolism and excretion. Consistent with such regulation is endothelial PIEZO1-dependence of hepatic, intestinal and whole body cholesterol and bile homeostasis. There is organ perfusion-dependent gene regulation via endothelial PIEZO1 and downstream nitric oxide synthase. Endothelial PIEZO1-deleted mice are protected against hyperlipidaemia and ectopic fat deposition. Human *PIEZO1* gene variants and a recapitulated human *PIEZO1* gain-of-function variant in mice associate with dyslipidaemia. The data suggest lipid-promoting effects of endothelial force sensing and new opportunity for understanding and addressing problems of hyperlipidaemia.

## INTRODUCTION

Cardiovascular disease was the leading cause of death in 2000 and 2019 and responsible for the largest increase in deaths worldwide over the last two decades ^1^. A major step in the fight against it has been recognition of the importance of dysregulated lipid homeostasis as a risk factor, leading to successful protective strategies such as statin medication to lower plasma cholesterol ^2^. The problem is nevertheless unsolved and remains a global challenge ^3^. A current focus of research is on hypertriglyceridemia with the suggestion of a new era of therapies aimed at lowering triglycerides to protect against cardiovascular and other diseases such as non-alcoholic fatty liver disease and pancreatitis ^3,4^. The interruption of sedentary behaviour (e.g., due to sitting down) by physical activity (e.g., light-intensity walking) attenuates post-prandial triglyceride elevation ^5^. A potential mechanism for such beneficial effects is the detection of mechanical force arising from cardiovascular activity, an important example of which is shear stress caused by blood flow against endothelium ^6,7^.

PIEZO proteins are special force sensors in eukaryotes ^8-11^. One of them, PIEZO1, is increasingly recognised as a key detector of shear stress at the endothelium ^12-24^. PIEZO1s associate in trimers to form large curved ion channels of 114 membrane-spanning segments and almost a million Daltons each, acting as sophisticated molecular machines that almost instantaneously couple mechanical force to transmembrane ion flux, thereby regulating cell function ^10,25-28^. Endothelial PIEZO1 has known importance in vascular development and structure ^14^, exercise-dependent blood pressure regulation ^16^, atherosclerosis ^18^, angiogenesis ^14,29^, vascular permeability ^30,31^, lymphatic valve formation ^32^, leukocyte diapedesis ^33^, muscle capillary density ^34^ and physical activity performance ^34^. It is notable also for its powerful positive regulation of one of the most important cardiovascular and metabolic mediators, nitric oxide ^14,19,24,35^.

Expression and function of PIEZO1 occur in endothelial cells of the liver ^13,14,16,36,37^, the major organ of lipid regulation ^38^. We therefore tested if endothelial PIEZO1 affects liver and systemic lipid homeostasis, beginning with unbiased analysis of gene expression in the liver.

## RESULTS

### Endothelial PIEZO1 inhibits *Cyp7a1* expression

Mice genetically engineered for conditional endothelial-specific deletion of PIEZO1 at the adult stage ^13,16^ were studied. Because lipid homeostasis depends on diet, we compared matched non-deleted PIEZO1 expressing mice (Control mice) with endothelial PIEZO1-deleted mice (PIEZO1^∆EC^ mice) in conditions of low fat diet (chow diet, CD) and high fat diet (HFD) comprising 60% fat. RNA sequencing was used to compare liver transcriptomes of these mice. Many potential differences are evident but only one is independent of dietary fat content. This unique change is an increase in *Cyp7a1* expression (Figure 1a, b). Independent quantitative PCR analysis validated the effect (Figure 1c). *Cyp7a1* encodes CP7A1 (Cytochrome P450 7A1), a key enzyme in cholesterol catabolism and classical bile acid synthesis, which accounts for a large fraction of bile ^39^. Control of this mechanism is largely transcriptional ^40^, so changes in *Cyp7a1* mRNA are likely to be functionally important. The data suggest a novel hypothesis in which mechanical force acting at endothelial PIEZO1 constitutively inhibits the conversion of cholesterol to bile by inhibiting *Cyp7a1*.

**Figure 1.**
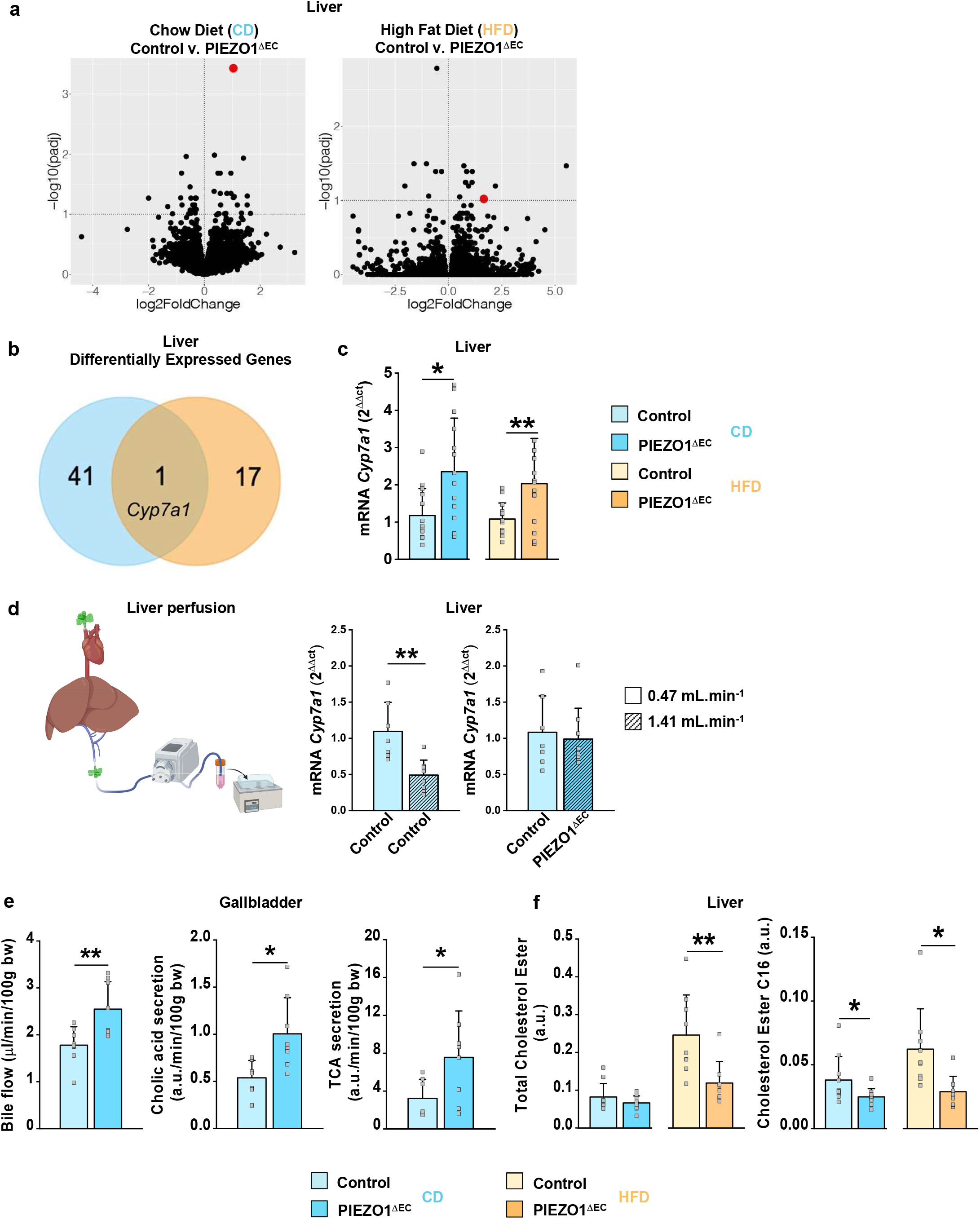
Regulation of *Cyp7a1* and associated hepatobiliary lipid homeostasis by endothelial PIEZO1. Measurements were made from 20 week-old mice fed chow diet (CD) and then CD (data in blue) or high fat diet (HFD, data in orange) for 8 weeks. Light coloured data indicate data from control (PIEZO1-expressing) mice and darker coloured data indicate data from endothelial PIEZO1-deleted (PIEZO1^∆EC^) mice. **a**. Volcano plots showing the adjusted P-values and the log2 fold change (FC) values of genes in the two genotype models (control mice versus (v.) PIEZO1^∆EC^ mice) fed CD (left) or HFD (right) (n = 5 per group). Differentially expressed genes (DEGs) were identified as genes with an adjusted p-value of < 0.1. The red dot in both data sets is for *Cyp7a1*. All data are deposited in ArrayExpress: <u>E-MTAB-9996</u>. **b**. Venn diagram representing all significant DEGs in the two dietary conditions (adjusted p-value of < 0.1). There were 41 DEGs unique to the CD condition (blue circle) and 17 unique to the HFD condition (orange circle). 1 DEG was common to both conditions and this was *Cyp7a1*. **c**. mRNA expression assessed by Q-PCR for *Cyp7a1* in liver of control and PIEZO1^∆EC^ mice fed CD or HFD (CD n = 14 per group; HFD n = 15 per group). mRNA expression was normalised to *rps19* mRNA abundance. **d**. On the left is a diagram illustrating the experimental arrangement for liver perfusion. On the right are bar charts showing liver *Cyp7a1* Q-PCR data from liver perfusion experiments at two perfusion rates for control mice (light blue, 0.47 mL.min^-1^, n = 10 mice; dark blue hatched, 1.41 mL.min^-1^, n = 10 mice) and PIEZO1^∆EC^ mice at 0.47 mL.min^-1^ (n = 10) and at 1.41 mL.min^-1^ (n = 9). **e**. Data shown as bar charts for gallbladder cannulation experiments comparing control mice and PIEZO1^∆EC^ mice on chow diet (CD) for bile flow (n = 8 mice for control, n = 8 mice for PIEZO1^∆EC^) and cholic acid and taurocholic acid (TCA) (n = 7 mice for control, n = 8 mice for PIEZO1^∆EC^) relative to body weight (bw). Cholic acid and TCA levels were expressed as arbitrary unit (a.u.). **f**. Relative quantity of cholesterol ester in liver shown as arbitrary unit (a.u.) (CD n = 10 control mice and n = 10 PIEZO1^∆EC^ mice; HFD n = 9 control mice and n = 8 PIEZO1^∆EC^ mice). On the right relative quantity of Cholesterol ester C16 (CD n = 10 mice per group; HFD n = 9 mice per group). Summary data are mean ± s.d.. Superimposed are the underlying individual data as open symbols. Statistically significant differences are indicated by: \**P*<0.05; \*\**P*<0.01.

### Portal flow dynamically regulates *Cyp7a1* via PIEZO1

*In vivo*, endothelial PIEZO1 channels are expected to be constitutively stimulated by fluid flow, so phenotypes resulting from PIEZO1 deletion are suggestive of an underlying mechanical effect. To test directly if mechanical force negatively regulates *Cyp7a1* expression and determine this gene’s susceptibility to physiological change in fluid flow, we cannulated mice to artificially control portal flow, which accounts for about 75% of the liver’s blood flow ^41^. In mice, portal flow is 1.6-2.3 mL.min^-1^ in basal conditions ^42^ but the flow is expected to vary from such values, for example decreasing during whole body physical activity ^43^. We therefore compared the effects of 0.47 and 1.41 mL.min^-1^ flow rates, firstly in control PIEZO1-expressing mice. There is less *Cyp7a1* expression at the high (1.41 mL.min^-1^) flow rate (Figure 1d). The effect of high flow is absent when PIEZO1 is deleted (Figure 1d). The data suggest that mechanical force generated by flow inhibits *Cyp7a1* expression and that it does so *via* endothelial PIEZO1.

### Endothelial PIEZO1 reduces bile synthesis and flow

Changes in *Cyp7a1* are expected to affect bile homeostasis ^39,40^. To test if this is the case, we performed gallbladder cannulation to measure bile flow and composition. Bile flow is increased in PIEZO1^∆EC^ mice (Figure 1e), consistent with *Cyp7a1* expression being upregulated by endothelial PIEZO1 deletion (Figure 1c). Accordingly, there are increased secretion rates of bile acid and the bile acid conjugates, cholic acid and taurocholic acid (TCA) ^44^ (Figure 1e). The data support the idea that stimulation of endothelial PIEZO1 by blood flow reduces bile flow by reducing hepatic *Cyp7a1* expression and therefore bile synthesis in the liver.

### Endothelial PIEZO1 increases liver cholesterol

Bile acids are produced by CP7A1-mediated cholesterol catalysis ^39,40^. Therefore, although there are many mechanisms controlling cholesterol, less cholesterol is expected when there is more bile synthesis. In PIEZO1^∆EC^ mice (which have increased bile acid secretion), there is less total cholesterol ester in liver in the high fat diet condition (Figure 1f). More detailed analysis revealed less cholesteryl palmitate (C16), a cholesterol ester obtained by condensation of cholesterol with palmitic acid, in chow and high fat diet conditions (Figure 1f), indicating differential effects on cholesterol subspecies. The data suggest that endothelial PIEZO1 increases cholesterol in the liver, consistent with it inhibiting *Cyp7a1*.

### Nitric oxide links endothelial PIEZO1 to *Cyp7a1*

Because the PIEZO1 deletion in our experiments is endothelial-specific ^16^ and *Cyp7a1* regulation occurs in hepatocytes ^45^, we speculated that these physically separate mechanisms are connected by a diffusible messenger. Prior studies show that PIEZO1 is a driver of nitric oxide production by endothelium ^14,19,24^ and that nitric oxide inhibits *Cyp7a1* expression ^45,46^. Therefore, we hypothesized interconnection via nitric oxide, which is readily diffusible ^35^. To test this idea, we isolated liver microvascular endothelial cells and biochemically investigated endothelial nitric oxide synthase (NOS3), which is phosphorylated at serine-1179 after PIEZO1 activation in other endothelial cell types ^14,19^. Such phosphorylation is commonly associated with enhanced NOS3 activity ^35^. In liver microvascular endothelial cells, PIEZO1 small-molecule agonist, Yoda1, significantly increases phosphorylation at serine-1179 and this effect depends absolutely on endothelial PIEZO1 (Figure 2a). NOS3 substrate inhibitor N^G^-methyl-L-arginine (L-NMMA) inhibits the effect of portal flow on *Cyp7a1* expression in wildtype mice (Figure 2b *cf* Figure 1d) and systemic administration of L-NMMA in freely moving wildtype mice selectively increases *Cyp7a1* expression (Figure 2c, SI Figure S1). The data suggest that nitric oxide links endothelial PIEZO1 to hepatocyte *Cyp7a1* (Figure 2d).

**Figure 2.**
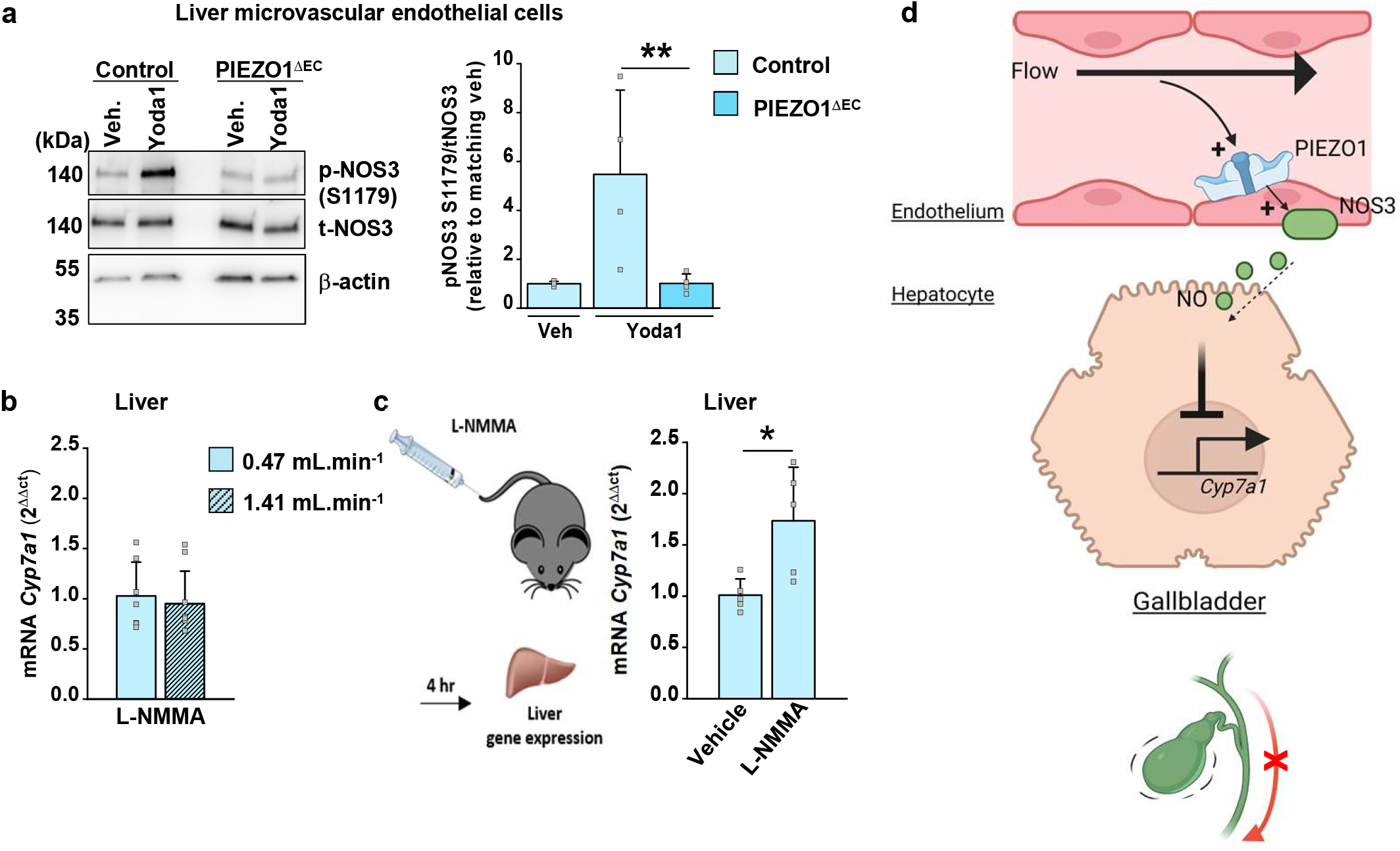
Nitric oxide links endothelial PIEZO1 to *Cyp7a1*. **a**. On the left, example Western blot for liver microvascular endothelial cell proteins from control and PIEZO1^∆EC^ mice after 1 min treatment with 2 µM Yoda1 or the vehicle (veh., DMSO) control. Blots were probed for NOS3 phosphorylation at S1179 (p-NOS3), total NOS3 (t-NOS3) and the loading control protein, β-actin. On the right, quantification for experiments of the type exemplified, expressed as phospho-NOS3 intensity relative to total NOS3 intensity and normalised to matching Vehicle condition (n = 4 each group). **b**. Bar chart showing liver *Cyp7a1* Q-PCR data from liver perfusion experiments in control mice of the type described in Figure 1d comparing the effects of the two perfusion rates, both with 0.3 mM L-NMMA in the perfusate (0.47 mL.min^-1^, n = 7 mice; hatched, 1.41 mL.min^-1^, n = 9 mice). **c**. Bar chart showing liver *Cyp7a1* Q-PCR data for wildtype mice fed CD and injected retro-orbitally with 72 µL of 1 mM L-NMMA per 100 g of body weight or vehicle control (PBS) (n = 5 each group). **d**. Schematic of the proposed signalling between endothelial PIEZO1 and hepatocyte *Cyp7a1*. Summary data are mean ± s.d.. Superimposed are the underlying individual data as open symbols. Statistically significant differences are indicated by: \**P*<0.05; \*\**P*<0.01.

### Endothelial PIEZO1 regulates lipids beyond the liver, cholesterol and bile

Cholesterol and bile regulations in the liver affect whole body (global) lipids ^40,44^. Therefore, we investigated beyond the hepatobiliary system, first measuring lipids in plasma. As in the gallbladder (Figure 1e), there are increases in plasma bile salts in PIEZO1^∆EC^ mice but the pattern of effects is now more complex and effects in the high fat diet condition are absent or different (Figure 3a *cf* Figure 1e). As in the liver (Figure 1f), plasma cholesterol species are less in PIEZO1^∆EC^ mice on high fat diet condition (Figure 3b). We measured other lipids too. Triglycerides and free fatty acids are less in the high fat diet condition (Figure 3c). The data suggest effects of endothelial PIEZO1 on cholesterol and bile beyond the liver and lipids associated with dyslipidaemia beyond cholesterol and bile acids.

**Figure 3.**
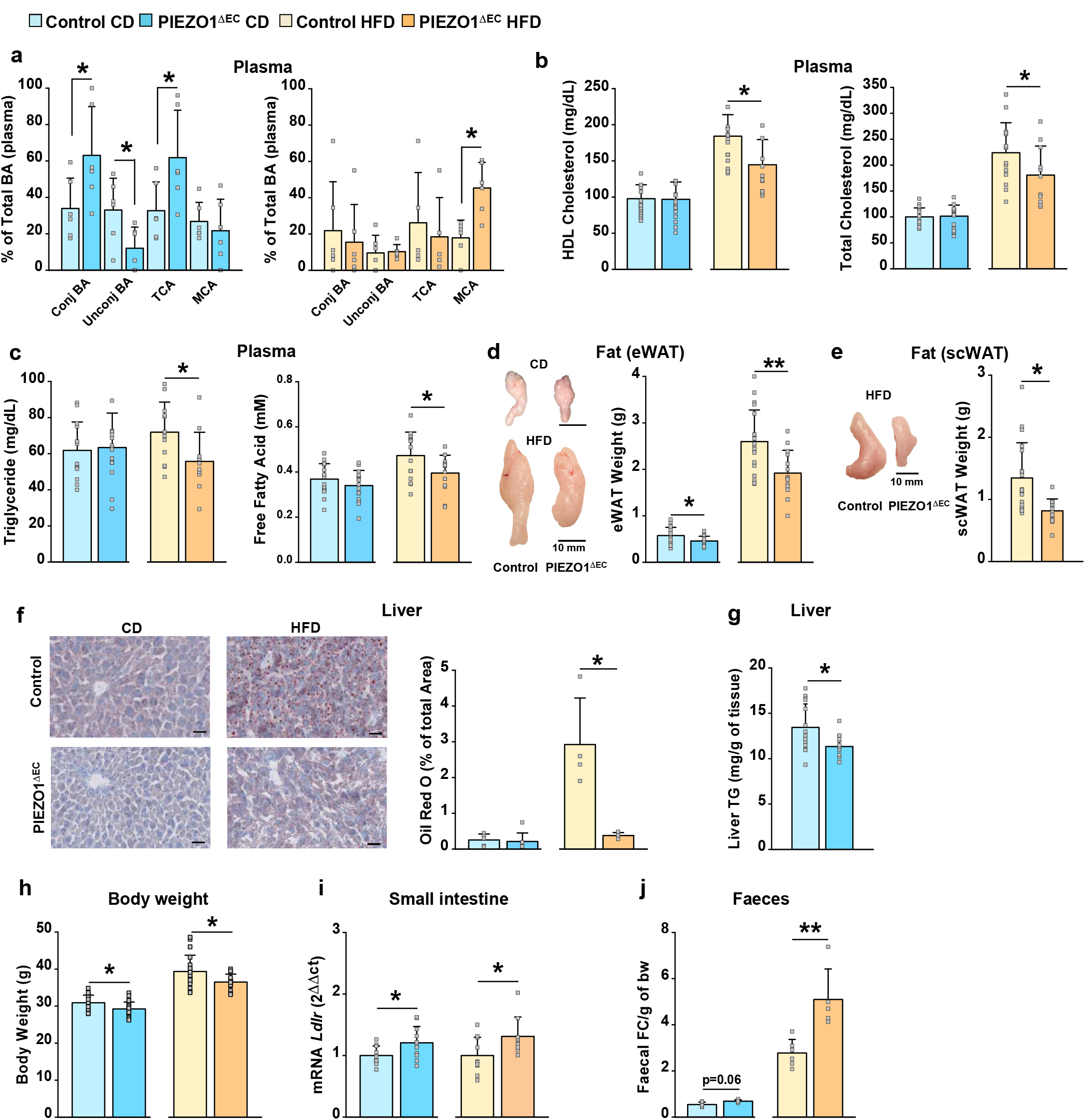
Endothelial PIEZO1 has global effects beyond *Cyp7a1*. **a**. Measurements were made from 20 week-old mice fed chow diet (CD) and then CD (data in blue) or high fat diet (HFD, data in orange) for 8 weeks. Light coloured data indicate data from control (PIEZO1-expressing) mice and darker coloured data indicate data from endothelial PIEZO1-deleted (PIEZO1^∆EC^) mice. **b**. Plasma conjugated and unconjugated CP7A1-regulated bile acid (BA), taurocholic acid (TCA) and muricholic acid (MCA) in plasma as percentages of total BA (n = 6 per group) **c**. Plasma concentrations of high-density lipoprotein (HDL) cholesterol (CD n = 17 control mice and n = 15 PIEZO1^∆EC^ mice; HFD n = 13 control mice and n = 11 PIEZO1^∆EC^ mice) and total cholesterol (CD n = 17 control mice and n = 16 PIEZO1^∆EC^ mice; HFD n = 14 control mice and n = 11 PIEZO1^∆EC^ mice). **c**. Plasma concentrations of triglycerides (CD n = 14 control mice and n = 15 PIEZO1^∆EC^ mice; HFD n = 12 control mice and n = 11 PIEZO1^∆EC^ mice) and free fatty acids (CD n = 17 control mice and n = 16 PIEZO1^∆EC^ mice; HFD n = 15 control mice and n = 14 PIEZO1^∆EC^ mice). **d**. On the left: Representative images of epididymal white adipose tissue (eWAT). Scale bars, 10 mm. On the right: Quantified weights for eWAT (CD n = 18 control mice n = 15 PIEZO1^∆EC^ mice; HFD n = 19 control mice n = 14 PIEZO1^∆EC^ mice). **e**. Representative images of Oil Red O-stained liver sections of control and PIEZO1^∆EC^ mice. Scale bars, 3 mm. On the right: Quantification of Oil Red O staining as exemplified on the left: n = 5 Control, n = 7 for PIEZO1^∆EC^. **f**. Hepatic triglyceride content (n = 15 Control mice, n = 13 PIEZO1^∆EC^ mice). **g**. Body weights (CD n = 23 per group; HFD n = 23 control mice and n = 19 PIEZO1^∆EC^ mice). **h**. Quantification of *Ldlr* mRNA abundance in distal small intestine (CD n = 10 control mice n = 11 PIEZO1^∆EC^ mice; HFD n = 10 control mice n = 9 PIEZO1^∆EC^ mice). **i**. Quantification of *Ldlr* mRNA abundance in distal small intestine of wildtype mice fed CD and injected retro-orbitally with 72 µL of 1 mM L-NMMA per 100 g of body weight or vehicle control (PBS) (n = 8 each group). **j**. Faecal free cholesterol (FC) per gram of body weight (n = 6 Control, n = 5 PIEZO1^∆EC^). Summary data are mean ± s.d.. Superimposed are the underlying individual data as open symbols. Statistically significant differences are indicated by: \**P*<0.05; \*\**P*<0.01.

### Endothelial PIEZO1 increases fat storage

Changes in plasma lipids are likely to have consequences for fat storage and so we measured fat deposits. Epididymal adipose tissue is smaller in PIEZO1^∆EC^ mice on chow and high fat diet (Figure 3d). In the high fat diet condition, there are lipid droplets in the liver (Figure 3e). These droplets are less visible in PIEZO1^∆EC^ mice (Figure 3e). Suspecting effects on liver lipid content in the chow diet condition also, we measured lipid triglycerides as a more sensitive indicator. There are lower liver triglycerides in PIEZO1^∆EC^ mice in the chow diet condition (Figure 3f). Consistent with the effects on fat storage, body weight is lower in PIEZO1^∆EC^ mice on chow and high fat diets (Figure 3g). The data support the idea that endothelial PIEZO1 affects whole body lipid homeostasis, increasing fat storage and body weight.

### Endothelial PIEZO1 decreases intestinal *Ldlr* expression and lipid excretion

Systemic lipid-lowering effects of deleting endothelial PIEZO1 are not readily explained by upregulation of bile, which solubilises intestinal lipids, increasing their availability for uptake ^39^. We therefore hypothesised an additional effect of endothelial PIEZO1 beyond the liver. Another major site of lipid control is the small intestine. The length of the small intestine is not different in PIEZO1^∆EC^ mice (SI Figure S2), so we quantified the expression of genes known to control lipid transport in proximal and distal small intestine (Figure 3h, SI Figure S2). We detected only one endothelial PIEZO1-related change in both diet conditions, which is increased *Ldlr* transcript in distal small intestine (Figure 3h). We hypothesized signalling via nitric oxide synthase, as we saw for *Cyp7a1* in liver (Figure 2). In support of this idea, systematic administration of L-NMMA selectively increases *Ldlr* expression in intestine (Figure 3i and SI Figure S2). Such an effect on *Ldlr* is relevant because it encodes LDLR protein (Low-density lipoprotein receptor). Specific overexpression of LDLR in intestine enhances cholesterol and total lipid excretion, and lowers body weight ^47^. We therefore measured lipid excretion. Free cholesterol in faeces is greater in PIEZO1^∆EC^ mice on high fat diet (Figure 3j). There is a potential similar effect in mice on chow diet (Figure 3j). Neutral lipids and free cholesterol are also increased in faeces of PIEZO1^∆EC^ mice (SI Figure S3). Effects in the high fat diet condition align with the results for plasma lipids (Figure 3b, c) but not fat storage, body fat or *Ldlr* transcript, which are affected similarly in both dietary conditions (Figure 3d-h). Technical limitations may have prevented detection of small changes in the chow diet condition (Figure 3e) but changes in the expression of other genes may also contribute (Figure 1a, SI Figure S2). The data suggest that an additional effect of endothelial PIEZO1 is to suppress *Ldlr* expression in the small intestine and decrease lipid excretion.

### Human *PIEZO1* variants associate with hepatobiliary phenotypes

To investigate if PIEZO1 is relevant to lipid homeostasis in humans, we interrogated UK Biobank and FinnGen databases using a candidate gene approach to determine if *PIEZO1* gene sequence variations associate with lipid disturbances or hepatobiliary disease. We find evidence of 14 *PIEZO1* variants associating (SI Table S1). Phenotypes range from death attributed to fatty liver, alcoholic liver disease, liver biliary pancreas problems, toxic liver disease and non-alcoholic fatty liver disease (SI Table S1). Moreover, the Cardiovascular Disease Knowledge Portal associates *PIEZO1* variants with LDL-cholesterol and total cholesterol (SI Table 1). Finally the NHGRI-EBI catalog of GWAS associates PIEZO1 variants with BMI-adjusted waist circumference (SI Table 1). Such data are not endothelial specific but suggest PIEZO1’s relevance in human lipid homeostasis.

### Global PIEZO1 gain-of-function disturbs lipid homeostasis

To determine potential implications of global changes in PIEZO1 (as may occur with *PIEZO1* variants), we took advantage of recapitulated human missense gain-of-function mutation in the germline of mice (PIEZO1^M-R/M-R^ mice), which amplifies PIEZO1 channel function and reproduces stomatocytosis found in people who carry such mutation ^48^. *Cyp7a1* expression, bile flow and cholic acid secretion are decreased and liver triglycerides are increased in these mice (Figure 4a-d). The data suggest that global gain-of-function PIEZO1 mutation has effects that mirror those of endothelial-specific PIEZO1 deletion, showing reduced bile secretion and increased liver triglycerides.

**Figure 4.**
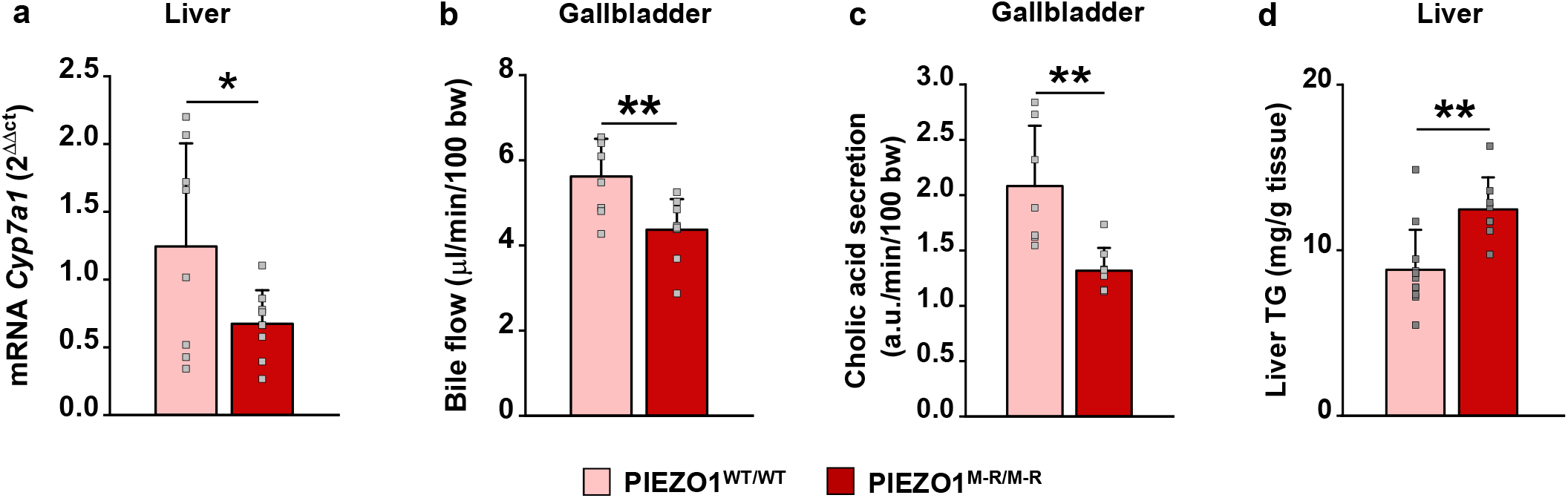
Global PIEZO1 gain-of-function mutation dysregulates lipids. Measurements were made from 8 week-old mice fed chow diet. Blue-coloured data indicate data from wildtype (PIEZO1^WT/WT^) control mice and light blue-coloured data indicate data from PIEZO1 gain-of-function mice (M-R homozygote, PIEZO1^M-R/M-R^). **a**. Liver *Cyp7a1* Q-PCR data (n = 8 PIEZO1^WT/WT^, n = 9 PIEZO1^M-R/M-R^). **b, c**. Bile flow data and LC-MS bile acid quantification from gallbladder cannulation experiments (Bile flow n = 8 PIEZO1^WT/WT^, n = 9 PIEZO1^M-R/M-R^; Cholic acid n = 8 PIEZO1^WT/WT^, n = 8 PIEZO1^M-R/M-R^). **d**. Hepatic triglyceride (TG) content (n = 12 PIEZO1^WT/WT^, n = 8 PIEZO1^M-R/M-R^) Summary data are mean ± s.d. Charts show the underlying individual data superimposed as open symbols. Significant differences: \**P*<0.05; \*\**P*<0.01.

## DISCUSSION

Based on these findings, we suggest control of global lipid homeostasis by mechanical force sensing. We propose a model in which microvascular endothelial cells of organs serve as force sensing units that use PIEZO1 to detect force from blood flow and signal its presence to surrounding cells by diffusible messengers such as nitric oxide (Figure 5). Pivotal genes of lipid biology are affected independently of dietary fat: *Cyp7a1*, which encodes the rate-limiting enzyme of cholesterol catabolism and bile acid synthesis, and *Ldlr*, which encodes a key receptor of cholesterol excretion. We suggest repression of these genes by the proposed force sensing mechanism with the common result of elevated systemic cholesterol. In many physiological settings, benefits are expected because cholesterol is an essential factor for membrane structure and steroid synthesis. We also suggest effects beyond cholesterol, more generally on other important lipids such as triglycerides.

**Figure 5.**
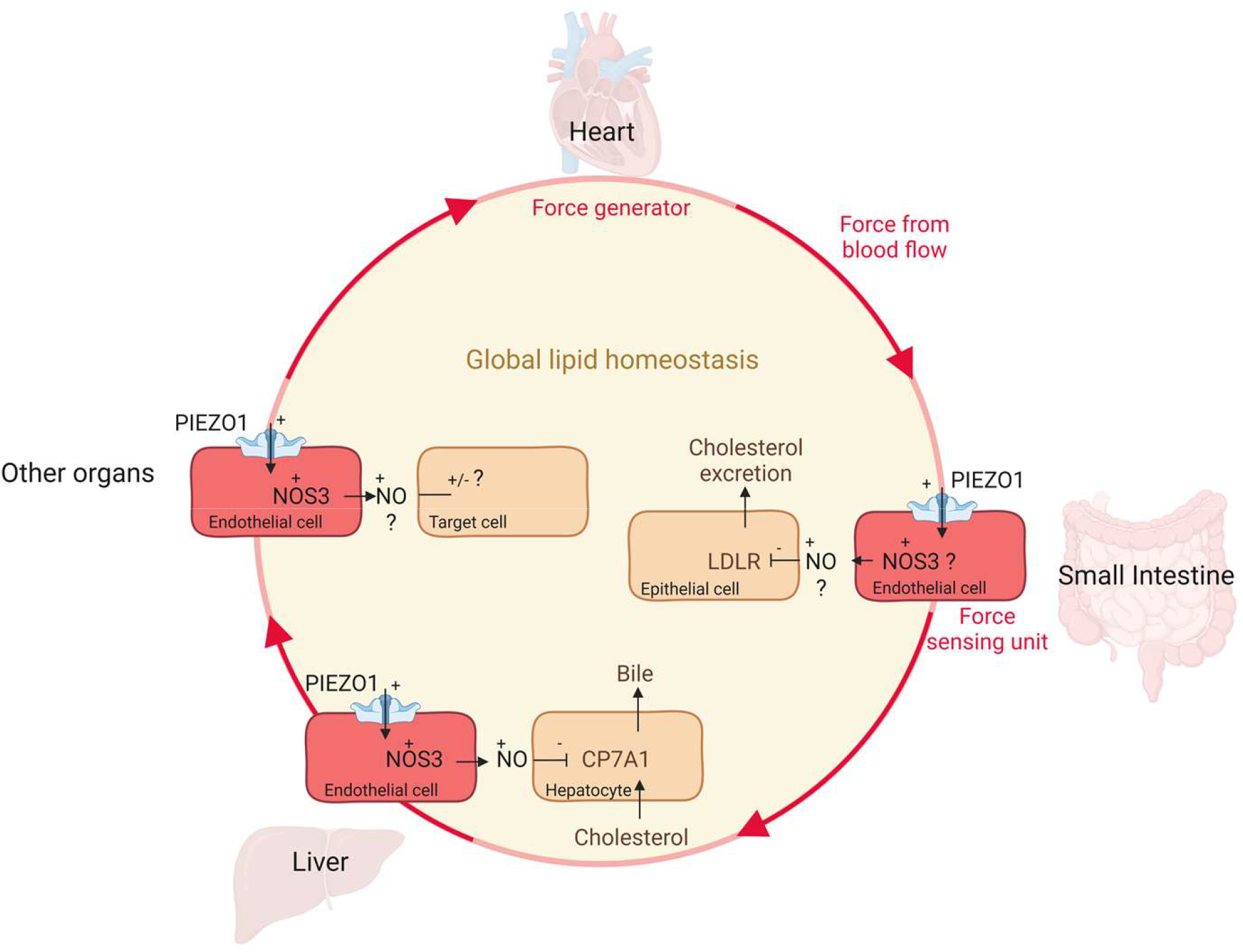
Model of how blood flow generates mechanical force to promote retention of cholesterol and other essential lipids. Shown is a schematic of the proposed mechanism in which the heart acts as the force generator, driving blood flow into microvasculature of organs such as the small intestine and liver. The endothelial cells act as blood flow sensors, in which PIEZO1 is a key mechanical detector. Once activated, PIEZO1 signals to downstream pathways such as NOS3 (also commonly referred to as eNOS) to generate diffusible nitric oxide (NO). NO is suggested to suppress the expression of key lipid regulatory genes such as *Ldlr*, which encodes LDLR, and *Cyp7a1*, which encodes CP7A1. Less LDLR leads in less cholesterol excretion and therefore more circulating cholesterol. Less CP7A1 results in less cholesterol catabolism to bile acids and also more circulating cholesterol. Similar mechanisms are suggested to exist in other organs, also regulating lipids. The overall effect is elevated cholesterol. Other lipids such as triglycerides are also elevated, through mechanism yet to be elucidated.

Our proposed mechanism of nitric oxide acting as a diffusible factor linking endothelium to surrounding cells is consistent with previous work showing an essential role of nitric oxide as an inhibitor of bile salt production in situations of bile salt excess ^45,46^. Nitric oxide has been shown to signal to the nucleus of hepatocytes via GAPDH ^45^. We expand this concept beyond local regulation in hepatocytes and bile excess to physiological regulation of bile salts by endothelial force sensing through PIEZO1 and nitric oxide diffusion to hepatocytes. Signalling of endothelial PIEZO1 via NOS3 is likely to be important in many blood vessels and organs because of the widespread expression of NOS3 in endothelium and diverse effects of nitric oxide ^35^. Nevertheless, the *Cyp7a1* and *Ldlr* effects are remarkably focussed on these genes; they were the only gene upregulations identified in low and high fat diet dietary conditions and following L-NMMA administration. How such selectivity arises is unknown. One possibility is exceptional nitric oxide sensitivity of these genes, but other factors may be involved. Endothelial PIEZO1 also signals, for example, through protease ADAM10 ^13^, the NOTCH1 pathway ^13^ and ATP ^19^. There might therefore be multi-layered signalling that ensures selective gene regulation.

We propose mechanical regulation of lipid metabolism via PIEZO1, a protein that is integral to membranes and itself lipid sensitive. PIEZO1 is inhibited by cholesterol and dietary saturated fatty acids ^49,50^ and stimulated or sensitized by phosphatidylinositol 4,5-bisphosphate ^49,51,52^, ceramide ^17^ and the cholesterol binding protein stomatin-like protein-3 ^53^. Lipids in the ion pore region of PIEZO1 channels are a feature of the closed channel state ^54,55^. Some of these lipid effects on PIEZO1 may be relevant to its regulation of lipids, such as the effect of cholesterol on PIEZO1, which may confer a negative feedback mechanism on PIEZO1 when systemic cholesterol increases.

The mice we studied were physically active but in a laboratory cage without access to a running wheel or similar device. They were therefore relatively sedentary. Our work raises the question of what would happen when physical activity levels increase. Physical activity changes blood flow distribution in an organ-specific manner, increasing blood flow in organs such as skeletal muscle but decreasing it elsewhere such as in the liver and intestines to prioritise blood for muscle activity ^7,43,56,57^. Therefore, physical activity is predicted to decrease cholesterol through the endothelial PIEZO1 mechanism we propose; i.e., because blood flow decreases in the liver and intestines in exercise, reducing nitric oxide and promoting *Cyp7a1* and *Ldlr* expressions. This would be consistent with the known lipid-lowering effects of physical exercise ^58^.

PIEZO1 is expressed in multiple cell types ^10^ and so may also have effects on lipid homeostasis via non-endothelial cells. Adipocyte PIEZO1 reduces adipocyte size and adipogenesis ^59,60^. PIEZO1 stimulates cholesterol biosynthesis in neural stem cells ^61^. PIEZO1 of cholangiocytes may regulate bile duct activity ^62^. There are also other ways in which PIEZO1 may act globally, such as through its functions in muscle and bone cells ^10,63-65^. These other roles of PIEZO1 could be important when there is gain-of-function mutation in PIEZO1, which is germline and global, as we describe here for PIEZO1^MR-MR^ mice. Nevertheless, some of the effects we see in PIEZO1^MR-MR^ mice are remarkably predictable from our endothelial PIEZO1 deletion studies (i.e., opposite to them), perhaps reflecting the importance of endothelial cells in all organs.

Gain-of-function PIEZO1 mutations are increasingly recognised and seen to be more frequent in some people, such as those of African, African American and South Asian ancestry ^66-68^. There are associations with anaemia, malarial resistance, iron overload, sprinting ability, itch, cardiac hypertrophy and HbA1c, a marker of diabetes ^66,68-72^. Here, we raise the possibility of additional associations with lipid homeostasis and hepatobiliary conditions such as fatty liver disease. Gain-of-function mutations in *PIEZO1* are suggested to have been selected evolutionarily to protect against malaria ^66,67^ but there may also have been other advantages such as through increased cholesterol, increased fat storage and reduced bile acid diarrhoea in times of cyclical food availability. In many modern human societies that have continuous and often excess food availability, fat-rich diets and sedentary lifestyles, disadvantages are likely through increased risk of poor digestion, obesity and cardiovascular and other diseases. It may therefore be worth considering PIEZO1 inhibition as a potential route to new therapeutics, for example to address dyslipidaemias such as hypertriglyceridemia ^3,4^ because we show that PIEZO1 deletion reduces plasma and liver triglycerides. Small-molecule modulators of PIEZO1 channels have been identified ^37,73-75^, so it may be possible to achieve suitable pharmacology and generate new therapeutics via further studies of this target.

## METHODS

### PIEZO1^∆EC^ mice

All animal use was authorized by the University of Leeds Animal Ethics Committee and The Home Office, UK. Animals were maintained in GM500 individually ventilated cages (Animal Care Systems) at 21 °C, 50–70% humidity, light/dark cycle 12/12 h on chow diet (CD) or 60% fat (HFD) (catalogue #F3282) ad libitum and bedding of Pure’o Cell (Special Diet Services, Datesand Ltd, Manchester, UK). Genotypes were determined using real-time PCR with specific probes designed for each gene (Transnetyx, Cordova, TN). C57BL/6 J mice with *Piezo1* gene flanked with LoxP sites (*Piezo1*-floxed) were described previously ^16^. To generate tamoxifen (TAM, Sigma-Aldrich) inducible disruption of *Piezo1* gene in endothelium, *Piezo1*-floxed mice were crossed with mice expressing cre recombinase under the endothelial-specific *Cadh5* promoter (Tg(*Cdh5*-cre/ERT2)1Rha and inbred to obtain *Piezo1*flox/flox/*Cdh5*-cre mice ^16^. TAM was dissolved in corn oil (Sigma-Aldrich) at 20 mg.mL^-1^. Mice were injected intra-peritoneally with 75 mg.kg^-1^ TAM for 5 consecutive days and studies were performed up to 10 weeks later. *Piezo1*flox/flox/*Cdh5*-cre mice that received TAM injections are referred to as PIEZO1^ΔEC^ mice. *Piezo1*flox/flox littermates (lacking *Cdh5*-cre) that received TAM injections were the controls (Control mice).

### PIEZO1^M-R/M-R^ mice

Mice were generated using CRISPR/Cas9 methodology. The guide RNA (gRNA) was designed to direct Cas9-mediated cleavage of Piezo1 6-bp upstream of the target methionine codon with no off-target sites of less than 3 mismatches predicted elsewhere in the genome (binding sequence 5’ GGGCGCUCAUGGUGAACAG 3’). A 120 nucleotide single-stranded homology-directed repair (ssHDR) template was designed to incorporate the methionine to arginine missense mutation, in addition to silent mutations introducing a Mlu1 restriction enzyme recognition site to facilitate genotyping (5’ccacccgtcccctgagcctg aggggctccatgctgagcgtgcttccatccccagccaTTAttTacGCGTagcgcccagcagccatccattgtgccattcacaccccaggcctacgaggag 3’; capital letters denote the mutated bases) (Sigma Aldrich). The gRNA, delivered as an alt-R crRNA combined with tracrRNA (Integrated DNA Technologies, Illnois, USA), Cas9 protein (New England Biolabs) and PAGE purified ssHDR template (Integrated DNA Technologies, Illinois, USA) were microinjected into C57BL/6 mouse zygotes and implanted in females (University of Manchester Transgenic unit). Successful gene editing of pups was identified by Mlu1 digestion of PCR amplicons 3 (F: 5’ TCTGGTTCCCTCTGCTCTTC 3’, R: 5’ TGCCTTCGTGCCGTACTG 3’) and confirmed by DNA sequencing following sub-cloning into pCRTM-Blunt (Invitrogen). Further genotypes were determined by real-time PCR with specific probes designed for each gene (Transnetyx, Cordova, TN). Eight week-old male mice were used for experiments unless otherwise stated. Only male mice were used for experiments.

### Collection of blood

Systemic blood was collected in EDTA coated tubes from the vena cava and centrifuged (4000×g, 4 °C, 10 min). Plasma was aliquoted and snap-frozen in liquid nitrogen. Organs were dissected, their weight was recorded, and they were either directly snap-frozen in liquid nitrogen or fixed in 4% formalin for 48 h. Frozen samples were stored at -80 °C until analysis.

### RNA extraction and real-time qPCR analysis

Total RNA was prepared from tissues using TriZol reagent (Sigma). To extract RNA, chloroform was added to the samples. After centrifugation, the aqueous phase was collected and isopropanol was added to precipitate the RNA. The RNA pellet was washed and suspended in RNase-free water. Reverse transcription of mRNA was performed using Superscript® III Reverse Transcriptase (Invitrogen) in the presence of random primers (Promega) and RNase OUT™ (Invitrogen). Briefly, 1 μg of total RNA from each sample was used with 2 μL of random Hexamer (20 μmol.L^-1^) and 0.5 μL of RNase OUT. After 8 min of incubation at 70 °C to inactivate DNase, 100 units of M-MLV transcriptase (Invitrogen) were added and the mixture was incubated at 75 °C for 7 min, 10 min at RT and then at 37 °C for 1 h. The RT reaction was ended by heating the mixture at 95 °C for 5 min. It was then chilled and stored at -80 °C until use. Real-time quantitative PCR analysis was performed using SYBR Green I Master (Roche) on LightCycler® 480 Real-Time PCR System. PCR reactions were run in a 384-well format with 10 μL reaction mixture containing SYBR Green I Master (Biorad) (5 μL), cDNA from the RT reaction (1 μL) and gene specific primers (1.25 µM, 4 μL). Reactions were run with a standard 2-step cycling program, 95 °C for 10 s and 60 °C for 40 s, for 40 cycles. Each analysis included both housekeeping gene and gene of interest, primer pairs as multiplexed samples. mRNA abundance was calculated relative to the average of the housekeeping gene rps19 and further normalized to the relative expression level of the respective controls (8-10 mice per group). The reactions were analysed with the ΔΔCT analysis method. PCR primers are specified in SI Table S2.

### RNA sequencing and data processing

The quality of RNA samples was assessed using Nanodrop, agarose gel electrophoresis and Agilent 2100. RNA sequencing (RNA-seq) was performed by Novogene Co., Ltd, Cambridge using Illumina Novaseq platform with 150-bp paired-ends. Raw data were filtered to remove low quality reads (Qscore <= 5) and adapters. Each sample generated more than 60 million clean reads. Raw reads in FASTQ format were provided by the sequencing facility. The initial quality control of the reads was performed using *FastQC* v0.118 ^76^. The reads were trimmed to remove low quality reads, using *TrimGalore* v0.6.2 ^77^ with a minimum quality score of 20 and at least length of 20 base pairs. Reads were aligned to the mouse reference genome (GRCm38 *Mus Musculus*) using *STAR* v2.7 ^78^ aligner. To improve the quality of the data, the alignments only retained uniquely mapped reads. Post-alignment quality control was performed. *featureCounts* v1.6 ^79^ was run with default parameters to quantify the read counts of each gene and used for differential expression analysis. The raw data are deposited in ArrayExpress: <u>E-MTAB-9996</u>.

### Differential expression analysis

Differential gene expression analysis was performed using the R-package *DESeq2* v1.26 ^80^. Low expression genes were removed from the analysis. Genes with a sum greater than ten read counts across all samples were retained in the analysis. *DESeq2* normalises the raw read counts based on the median of rations as described previously ^81^. Differentially expressed genes (DEGs) were identified with adjusted *P*-value below threshold 0.10 using the Benjamini-Hochberg correction procedure. Volcano plot and heat map of DEGs were constructed using the *ggplot2* package ^82^ and *pheatmap* function ^83^ implemented in *R*, respectively.

### In situ liver perfusion

8-12 week-old mice were then anaesthetized by intraperitoneal injection of ketamine hydrochloride (100 mg.kg^-1^) and xylazine hydrochloride (15 mg.kg^-1^). The portal vein was cannulated for inflow and the inferior vena cava for outflow. The liver was perfused for 5 min with PBS to flush out blood and prevent clotting of the hepatic vessels. Liver was then perfused with RPMI-1640 (Gibco 22400089) supplemented with 10% heat inactivated foetal bovine serum (FBS; Sigma, F9665-500ML) and 1% penicillin/streptomycin (Gibco) in a closed circuit for 2 hours with a peristaltic pump (Watson Marlow – 505Di). Flow was applied changing the flow rate of the peristaltic pump: 0.47 mL.min^-1^ or 1.41 mL.min^-1^.

### Liver sinusoidal endothelial cell isolation

Liver of 12-14 week-old male mice was used and tissue was mechanically separated using forceps, further cut in smaller pieces and incubated at 37 °C for 50 min, in a MACSMix Tube Rotator to provide continuous agitation, along with 0.1% Collagenase II (17101-015, Gibco, Waltham, MA) and Dispase Solution (17105-041, Gibco). Following enzymatic digestion samples were passed through 100 μm and 40 μm cell strainers to remove any undigested tissue. The suspension was incubated for 15 min with dead cell removal paramagnetic beads (130-090-101, Miltenyi Biotec GmbH, Bergisch Gladbach, Germany) and then passed through LS column (130-042-401, Miltenyi Biotec).

The cell suspension was incubated with CD146 magnetic beads (130-092-007, Miltenyi Biotec 130-092-007) at 4°C for 15 min under continuous agitation and passed through MS column (130-042-201, Miltenyi Biotec). The CD146 positive cells, retained in the MS column, were plunge out with PEB (PBS 1X, Bovine Serum Albumin 5 g.L^-1^, EDTA 2 mM) and centrifuged at 1000 RPM for 5 min. Cells were seeded on polyD-lysine coated 6 –well plates and cultured in EGM-2 media (C-22211, promocell Gmbh) supplemented with EGM2 supplement kit (C-39211, promocell Gmbh) until reaching 90% confluency. Cells were then starved in SBS for 1.5h prior to treatment with 2 µM Yoda1 or control vehicle (Veh., Dimethyl Sulfoxide DMSO) for 1 min. Cell pellet were lysed with NP40 lysis buffer (10 mM Tris pH7.4, 150 mM NaCl, 0.5 mM EDTA pH 8.0, 0.5% Nonidet P40, containing Phosphatase inhibitor (P0044, Sigma) and Protease inhibitor (P8340, Sigma).

### Gallbladder cannulation

Mice were then anaesthetized by intraperitoneal injection of ketamine hydrochloride (100 mg.kg^-1^) and xylazine hydrochloride (15 mg.kg^-1^). The common bile duct was ligated close to the duodenum; and then the gallbladder was punctured and cannulated with a polyethylene-10 catheter. Bile was collected for 30 min, after a stabilization time of 30 min. During bile collection, body temperature was stabilized using a temperature mattress. Bile flow (µL.min^-1^.100g of body weight (bw)^-1^) was determined gravimetrically assuming a density of 1 g.mL^-1^ for bile. Aliquots were snap frozen for later analysis.

### Immunoblotting

Protein concentration was determined using BCA assay and 20 µg of total protein were heated at 95 ̊C for 5 min in SDS-PAGE sample buffer, loaded on a precast 4– 20% polyacrylamide gradient gel (4561094 Biorad) and subjected to electrophoresis. Proteins were transferred onto a PVDF membrane using semi-dry transfer for 30 min. Membranes were blocked with 5% milk in Tris-buffered saline with Tween 0.05% for 1 hr at room temperature. The membranes were exposed to primary antibody overnight at 4 °C (Phospho eNOS S1179 -BD 612392; Total eNOS/NOS3: BD 610296; anti mouse: Jackson Immuno research 715-035-150; Beta Actin: Santa Cruz sc4778), rinsed and incubated with appropriate horseradish peroxidase-labelled secondary antibody for 1 hr at room temperature. The detection was performed by using SuperSignal West Femto (34096, ThermoFisher Scientific) and visualized with a G-Box Chemi-XT4 (SynGene, Cambridge, UK). Beta actin was used as reference protein.

### In vivo injection of L-NMMA

Mice of 20 weeks of aged were injected retro-orbitally with 10 mM L-NMMA (N^G^-methyl-L-arginine acetate salt, Tocris Bioscience, 0771) to reach a final systemic concentration of 10 µM. Liver and small intestine were harvested 4 hr later. mRNA was isolated as described above.

### LC-MS lipid quantification

A solution of 10 μM palmitoyl-L-carnitine-(N-methyl-d3) (Sigma), 10 μM palmitic acid-d31 (Sigma), and 10 μM deoxycholic acid-d6 (Sigma) in LC−MS-grade methanol was prepared as internal standard spiking solution (ISSS). Samples (from gall bladder cannulation) were reconstituted in 100 μL of LC−MS-grade water and 100 μL of ISSS, vortex-mixed, and sonicated for 30 min before being transferred to LC vials. Chromatography was performed using an Acquity UPLC system (Waters) equipped with a CORTECS T3 2.7 μm (2.1 × 30 mm) column, which was kept at 60 °C. The Acquity UPLC system was coupled to a Xevo TQ-XS mass spectrometer (Waters). The binary solvent system used was solvent A comprising LC−MS-grade water, 0.2 mM ammonium formate, and 0.01% formic acid and solvent B comprising analytical-grade acetonitrile/isopropanol 1:1, 0.2 mM ammonium formate, and 0.01% formic acid. For all analyses, a 10 μL injection was used, and the mobile phase was set at a flow rate of 1.3 mL.min^-1-^. For bile acid analysis, the column mobile phase was held at 20% solvent B for 0.1 min, followed by an increase from 20 to 55% solvent B over 0.7 min. The mobile phase was increased to 98% solvent B and held for 0.9 min. The mobile phase was then returned to 20% solvent B held for 0.1 min to re-equilibrate the column. Analyses were performed using multiple reaction monitoring (MRM).

### UK biobank, FinnGen, Cardiovascular Disease Knowledge Portal and NHGRI-EBI GAWS catalog

The chromosome 16 genomics data from UK Biobank was obtained under the identifier 42651. The data was filtered to excluded participants with more than 10% missing data, exclude variants with Hardy-Weinberg equilibrium (HWE) less than 1x10^-6^ and exclude variants with minor allele frequency (MAF) less than 5% for common variants or less than 1% for less frequent variants, using PLINK version 2.0 software. The related hepatobiliary diseases terms were obtained using the ID-20002 self-reported non-cancer illnesses code. Logistic regression analyses were performed using PLINK2 and adjusted for effects of sex, age and the first ten principal components. In FinnGen, we accessed the publicly available data using <u>FinnGenn r7</u>. Then, we screened for the PIEZO1 mutations linked to hepatobiliary diseases. This release contains 3,095 disease endpoints. CVD knowledge portal data are publicly available https://cvd.hugeamp.org/. EBI catalog is publicly available https://www.ebi.ac.uk/gwas/

### Measurement of neutral lipids

Lipids from 2 mg of liver or 100 mg of faeces were extracted according to previously described methods ^84^ in dichloromethane/methanol/water (2.5/2.5/2.1, v/v/v), in the presence of internal standards: 4 µg of stigmasterol, 4 µg of cholesteryl heptadecanoate, 8 µg of glyceryl trinonadecanoate. Dichloromethane phase was evaporated to dryness and dissolved in 30 µL of ethyl acetate. 1 µL of the lipid extract was analysed by gas-liquid chromatography on a FOCUS Thermo Electron system using a Zebron-1 Phenomenex fused silica capillary columns (5 m x 0.32 mm i.d. 0.50 µm film thickness) as previously described ^85^. Oven temperature was programmed from 200 °C to 350 °C at a rate of 5 °C per min and the carrier gas was hydrogen (0.5 bar). The injector and the detector were at 315°C and 345°C respectively.

### Measurement of plasma lipids

Total cholesterol and triglycerides (TGs) and HDL cholesterol concentrations were measured with commercial kits (CHOD-PAP for cholesterol #87656, GPO-PAP for TGs #87319 and Cholesterol HDL-PTA for HDL cholesterol #86516; BIOLABO at ABLIANCE, Compiègne, France). Non-esterified fatty acid (NEFA) concentrations were measured using a kit from WAKO chemicals.

### Measurement of plasma bile acids and salts

100 µL of plasma was diluted in 400 µL of ACN in the presence of an internal standard (23NorDCA, 20 ng). The mixture was agitated for 30 min at RT and centrifuged for 5 min at 10000 rpm. The supernatant was concentrated and reconstituted in 1 mL of 15 mM ammonium acetate (pH 5.3). The lipid extracts were pre-concentrated on HLB Solid-Phase Extraction (Waters SAS, Saint-Quentin-en-Yvelines). After conditioning of the plate (1 mL water, 2 mL methanol), samples were loaded and plates were washed with 2 mL of water and finally bile acids were eluted with 2 mL of methanol. After concentration, final extracts were solubilized in 20 μL of methanol and kept at -20°C until the analysis by LC-MS in high resolution. They were run on an Ultimate 3000 (ThermoFisher Scientific (Life Technologies SAS, Saint-Aubin, France) equipped with a ZORBAX SB-C18 column (2.1 x 100 mm, 1.7 μm, Agilent, Santa Clara, United States) kept at 45°C. Mobile phases were composed of (A) 15 mM ammonium acetate pH 5.3 (acetic acid), (B) acetonitrile and (C) methanol, in gradient mode at 0.5 mL.min^-1^. The gradient was as follow: 20% B/0%C at 0 min till 2.5min 16% B/20%C at 5 min; 1% B/95%C at 20 min; 1% B/95%C at 23 min; 20% B/1%C at 23.5 min till 27.5 min. The injection volume was 5 μL. High-resolution mass spectrometry detection was done on an Exactive (ThermoFisher Scientific, Life Technologies SAS, Saint-Aubin, France) in a ESI negative mode and an acquisition in full scan mode from 200 to 600 m/z. Parameter of the source were as follows: Sheath Gaz : 30 psi, Auxiliary Gas Flow : 10 Psi, Sweep as Flow Rate : 0 Psi, Spray voltage : 2500 V, Capillary Temperature : 200 °C, Capillary Voltage : -37.5 °C, Tube Lens Voltage : -165 V, Heater Temperature : 200 °C, Skimmer Voltage : -46 V, Maximum Injection Time : 250 ms. Calibration curves containing all the pure standards in methanol were run at different concentration (24, 6, 1.5, 0.375, 0.093, 0.023 and 0.0058 ng.mL^-1^). Quantification was carried out using the ion chromatogram obtained for each compound using 5 ppm windows with Trace Finder Quantitative Soft Ware (ThermoFisher Scientific -Life Technologies SAS, Saint-Aubin, France). The methodology was adapted from a previously described approach ^86^.

### Tissue staining

Liver tissues were frozen in O.C.T.™ compound (Tissue-Tek), 10-μm frozen liver sections were air-dried for 30 min and fixed in 4% formaldehyde. Oil Red O staining was performed using standard protocols. Tissue slides were scanned using an Axio Scan.Z1 on the Bioimaging platform at the University of Leeds. Oil Red O staining was quantified using Image J software.

### Statistical analysis

Genotypes of mice were always blinded to the experimenter and mice were studied in random order determined by the genotype of litters. Data sets comparing control and PIEZO1^ΔEC^ mice were analysed for statistical significance by independent student *t*-test without assuming equal variance. Outliers were removed in the validation analysis using the Robust regression and Outlier removal Test (ROUT) method with Q = 1% in GraphPad Prism 9.0 software. Statistical significance was considered to exist at probability *P* * < 0.05, ** < 0.01 or *** < 0.001. When data comparisons are shown without an asterisk, there were no significantly differences. The number of independent experiments is indicated by n. Descriptive statistics are shown as mean ± standard deviation (s.d.) unless indicated otherwise. All underlying individual data points are shown superimposed on mean data. Origin Pro software was used for data analysis and presentation unless otherwise indicated.

## DATA AVAILABILITY

A Source Data file is provided.

## ACKNOWLEDGEMENTS

The work was supported by research grants from Wellcome (grant number 110044/Z/15/Z) and British Heart Foundation (BHF) (RG/17/11/33042). LL and CWC were supported by BHF Mautner Fellowships and HJG and ELE were supported by BHF PhD Studentships.

For the purpose of Open Access, the authors have applied a CC BY public copyright license to any Author Accepted Manuscript version arising from this submission.

The UK Biobank data was accessed *via* application ID 42651 awarded to Vijayalakshmi Deivasikamani, who was a BHF Daphne Jackson Fellow.

Lipid profiling was performed at the Metatoul-lipidomic platform (INSERM UMR1048, Toulouse, France (MetaboHUB-ANR-11-INBS-0010) certified to ISO 9001:2015 standards. We thank Romana Mughal and Asjad Visnagri for technical assistance in animal studies.

## AUTHOR CONTRIBUTIONS

LL planned and coordinated experimental work and data analysis, proposed ideas for experiments, performed experiments, interpreted data and co-wrote the paper. CWC performed RNAseq data analysis and interpretation and wrote associated parts of the paper. ELE, HJG, FB, TSF and MR performed experimental studies and GP provided technical assistance. JBM provided lipidomic expertise. FB, ECB, SP, CK, GP, ELE, CWC, PS and LDR provided intellectual input. DJB initiated the project, generated research funds and ideas, led and coordinated the project, interpreted data and co-wrote the paper.

