## Supplementary material for "PIEZO1 force sensing controls global lipid homeostasis": Suplemental Information

### SUPPLEMENTAL INFORMATION (SI)

**SI Figure S1 Selectivity of systemic L-NMMA for *Cyp7a1***
Q-PCR mRNA expression data for 13 genes in liver of wildtype mice injected with 10 μM L-NMMA or vehicle control as described for Figure 2c. For each gene, L-NMMA data are shown as the fold-change relative its respective vehicle control data, presented as mean ± s.d. with underlying individual data for each mouse superimposed as open symbols. There were no statistically significant differences between L-NMMA and vehicle groups.

**SI Figure S2 Small intestine length and gene expression data**
Measurements were made from 20 week-old mice fed chow diet (CD) and then CD (data in blue) or high fat diet (HFD, data in orange) for 8 weeks. Light coloured data indicate data from control (PIEZO1-expressing) mice and darker coloured data indicate data from endothelial PIEZO1-deleted (PIEZO1$^{\Delta EC}$) mice.
**a**. Small intestine length (CD n = 10 control mice n = 8 PIEZO1$^{\Delta EC}$ mice; HFD n = 5 control mice n = 7 PIEZO1$^{\Delta EC}$ mice).
**b**, **c**. Q-PCR mRNA expression data for 12 genes in proximal (**b**) or distal (**c**) small intestine (CD n = 10 control mice n = 11 PIEZO1$^{\Delta EC}$ mice; HFD n = 11 control mice n = 10 PIEZO1$^{\Delta EC}$ mice).
**d**. Q-PCR mRNA expression data for additional genes in distal small intestine of wildtype mice injected with 10 μM L-NMMA or vehicle control as described for Figure 3i.
Summary data are mean ± s.d.. Superimposed are the underlying individual data as open symbols. Statistically significant differences are indicated by: *$P$<0.05; **$P$<0.01.

**SI Figure S3 Additional faecal lipid data**
Measurements were made from 20 week-old mice fed chow diet (CD) and then CD (data in blue) or high fat diet (HFD, data in orange) for 8 weeks. Light coloured data indicate data from control (PIEZO1-expressing) mice and darker coloured data indicate data from endothelial PIEZO1-deleted (PIEZO1$^{\Delta EC}$) mice.
**a**, **b**. Quantification of faecal neutral lipids (**a**) and free cholesterol (FC) (n = 6 Control groups, n = 5 PIEZO1$^{\Delta EC}$ groups) (**b**) (CD n = 6 control mice n = 5 PIEZO1$^{\Delta EC}$ mice; HFD n = 5 control mice n = 4 PIEZO1$^{\Delta EC}$ mice)
Summary data are mean ± s.d.. Superimposed are the underlying individual data as open symbols. Statistically significant differences are indicated by: *$P$<0.05.

**SI Figure S4 Uncropped western blot images for Figure 2a**

**SI Table S1 Human *PIEZO1* variants associated with hepatobiliary phenotypes**
P-values for *PIEZO1* single nucleotide polymorphism (SNP) associations with hepatobiliary phenotypes in people from UK Biobank, FinnGen, Cardiovascular Disease Knowledge Portal and NHGRI-EBI catalog.

**SI Table S2 Nucleotide sequences of Q-PCR primers used in this study**

### SI Figure S1

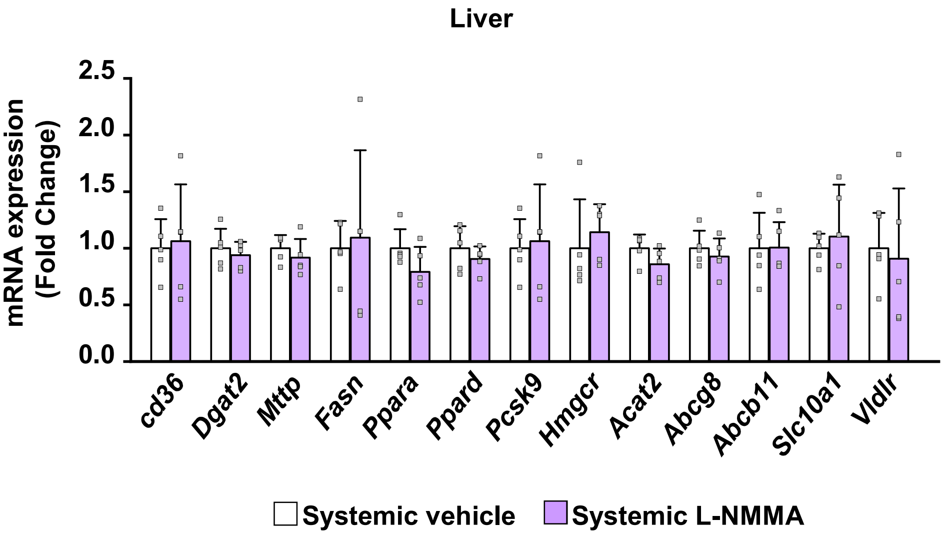
Liver
mRNA expression (Fold Change)
2.5
2.0
1.5
1.0
0.5
0.0
cd36
Dgat2
Mttp
Fasn
Ppara
Ppard
Pcsk9
Hmgcr
Acat2
Abcg8
Abcb11
Slc10a1
Vldlr
Systemic vehicle
Systemic L-NMMA

### SI Figure S2

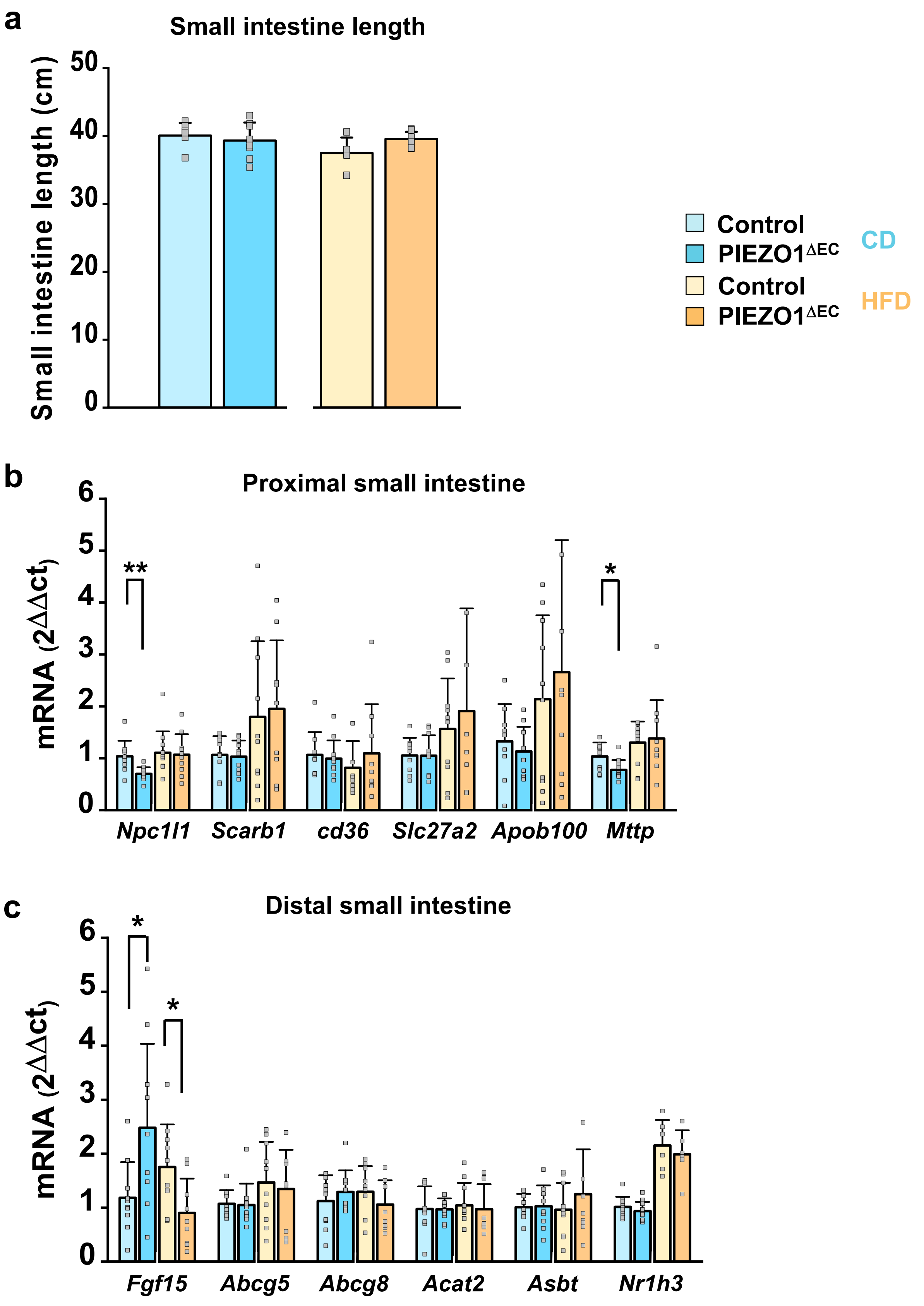

a
Small intestine length
Small intestine length (cm)
50
40
30
20
10
0
Control
PIEZO1ΔEC
Control
PIEZO1ΔEC
CD
HFD
b
Proximal small intestine
mRNA (2ΔΔct)
6
5
4
3
2
1
0
**
*
Npc1l1
Scarb1
cd36
Slc27a2
Apob100
Mttp
c
Distal small intestine
mRNA (2ΔΔct)
6
5
4
3
2
1
0
*
*
Fgf15
Abcg5
Abcg8
Acat2
Asbt
Nr1h3

#### SI Figure S3

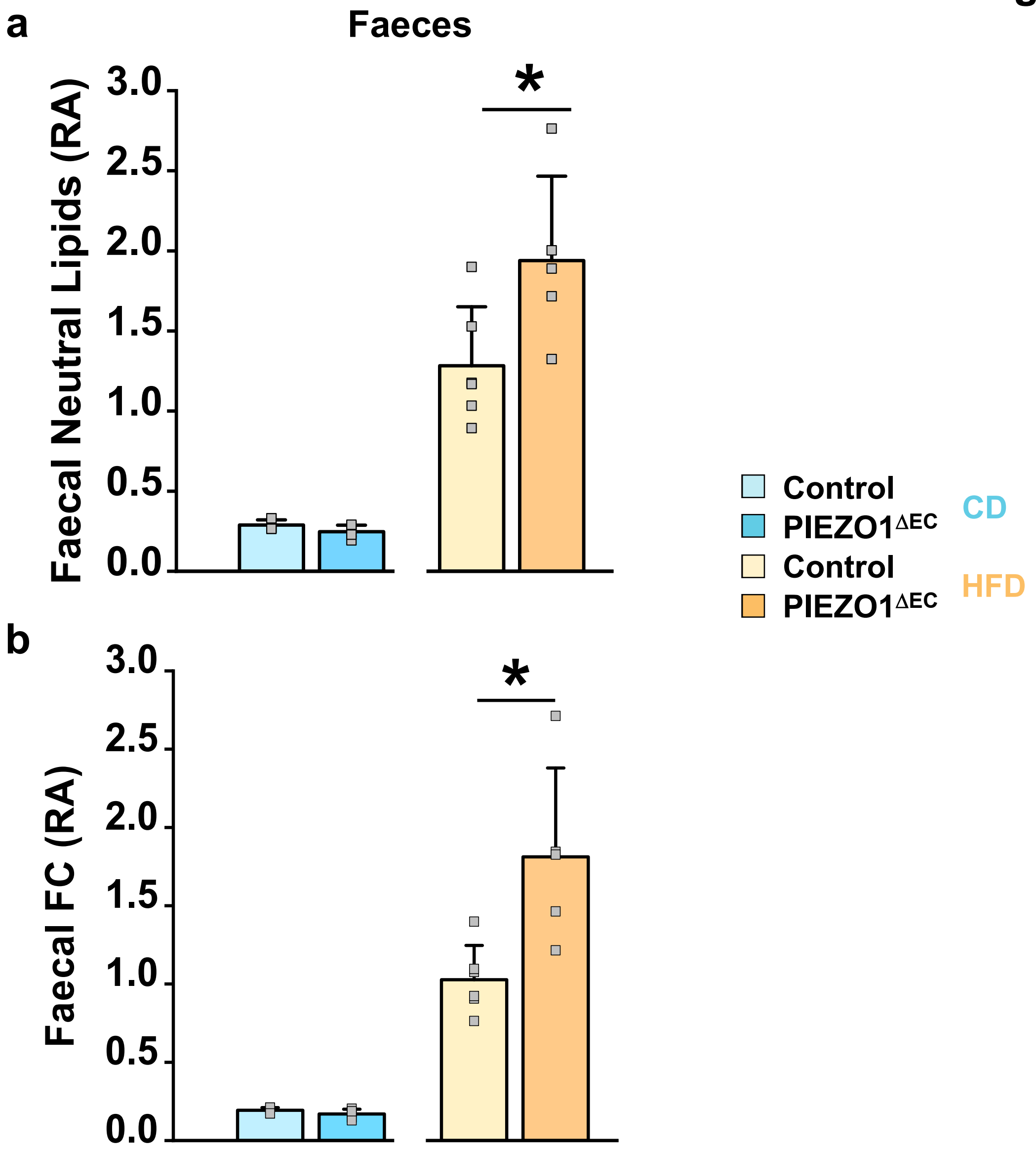
a
Faeces
Faecal Neutral Lipids (RA)
3.0
2.5
2.0
1.5
1.0
0.5
0.0
*
b
Faecal FC (RA)
3.0
2.5
2.0
1.5
1.0
0.5
0.0
*
Control
PIEZO1ΔEC
CD
Control
PIEZO1ΔEC
HFD

#### SI Figure S4

##### Uncropped western blots

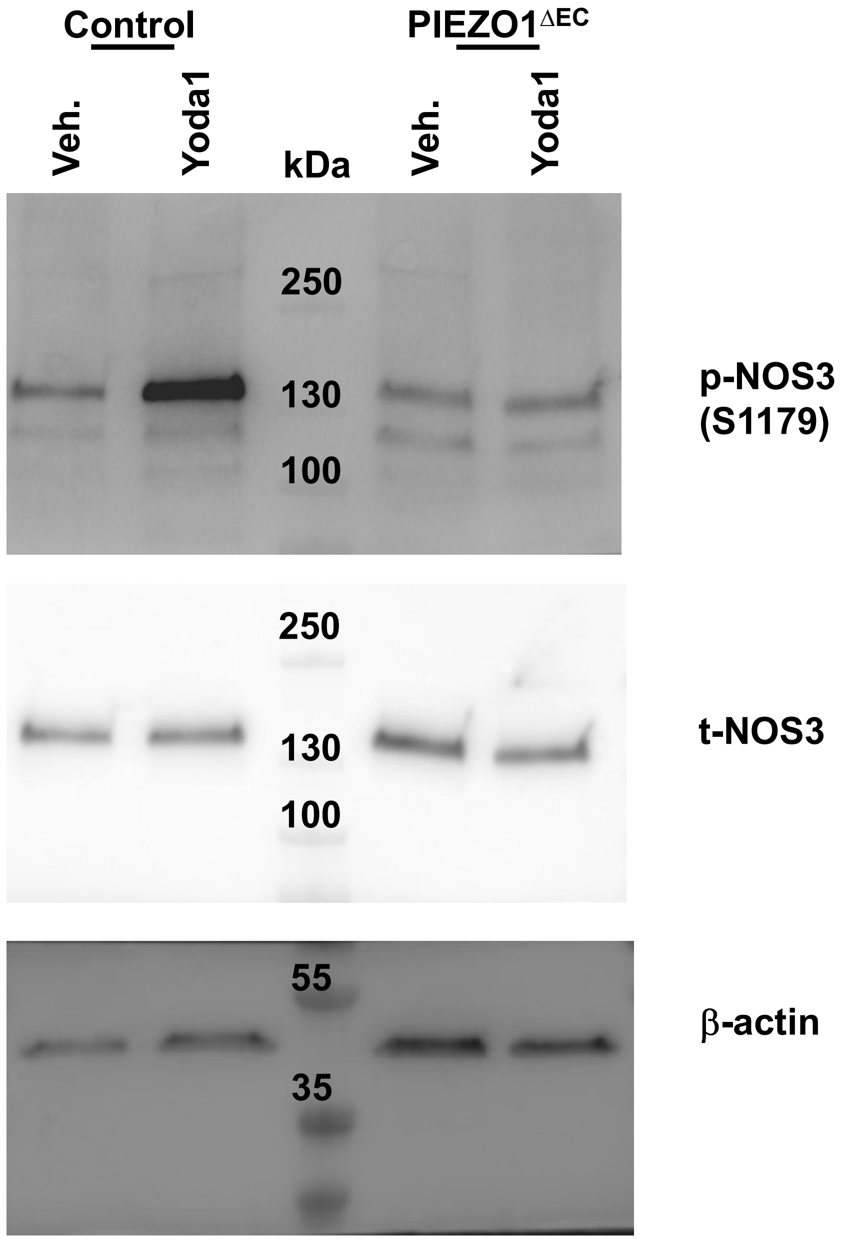

### SI Table S1

| rsid | *P*-value | Phenotype | Database |
|---|---|---|---|
| rs565024125 | 1.26E-14 | Cause of death: fatty liver | UK Biobank |
| rs377011269 | 2.21E-05 | Alcoholic liver diseases | UK Biobank |
| rs759885882 | 4.29E-07 | Alcoholic liver diseases | UK Biobank |
| rs115486714 | 6.85E-08 | Alcoholic liver diseases | UK Biobank |
| rs915624083 | 3.18E-05 | Liver biliary pancreas problem | UK Biobank |
| rs1044492299 | 1.65E-06 | Liver biliary pancreas problem | UK Biobank |
| rs95793364 | 9.78E-06 | Liver biliary pancreas problem | UK Biobank |
| rs747295957 | 3.18E-05 | Liver biliary pancreas problem | UK Biobank |
| rs942117829 | 6.35E-06 | Liver biliary pancreas problem | UK Biobank |
| rs201091400 | 4.74E-06 | Liver biliary pancreas problem | UK Biobank |
| rs1908139729 | 9.57E-06 | Liver biliary pancreas problem | UK Biobank |
| rs907173644 | 2.85E-05 | Liver biliary pancreas problem | UK Biobank |
| rs6500500 | 1.80E-04 | Toxic liver disease | FinnGen |
| rs561783880 | 7.60E-04 | Non-alcoholic fatty liver disease | FinnGen |
| rs9933309 | 5.14E-07 | LDL cholesterol | CVDKP |
| rs2608604 | 6.39E-07 | LDL cholesterol | CVDKP |
| rs2932690 | 3.34E-06 | LDL cholesterol | CVDKP |
| rs2911463 | 3.92E-06 | LDL cholesterol | CVDKP |
| rs9932423 | 7.24E-06 | LDL cholesterol | CVDKP |
| rs2002833 | 1.44E-05 | LDL cholesterol | CVDKP |
| rs8052231 | 1.81E-05 | LDL cholesterol | CVDKP |
| rs2911460 | 1.91E-05 | LDL cholesterol | CVDKP |
| rs8052370 | 3.04E-05 | LDL cholesterol | CVDKP |
| rs2926772 | 5.98E-05 | LDL cholesterol | CVDKP |
| rs2932690 | 6.15E-08 | Total cholesterol | CVDKP |
| rs2608604 | 9.36E-08 | Total cholesterol | CVDKP |
| rs8052231 | 9.57E-08 | Total cholesterol | CVDKP |
| rs9932423 | 2.02E-07 | Total cholesterol | CVDKP |
| rs8052370 | 4.42E-07 | Total cholesterol | CVDKP |
| rs9933309 | 5.17E-07 | Total cholesterol | CVDKP |
| rs2002833 | 7.47E-07 | Total cholesterol | CVDKP |
| rs2911460 | 1.15E-06 | Total cholesterol | CVDKP |
| rs57953994 | 3.17E-06 | Total cholesterol | CVDKP |
| rs475596 | 4.27E-06 | Total cholesterol | CVDKP |
| rs10445033 | 8.42E-06 | Total cholesterol | CVDKP |
| rs889764 | 1.21E-05 | Total cholesterol | CVDKP |
| rs6500505 | 1.22E-05 | Total cholesterol | CVDKP |
| rs2911463 | 1.38E-05 | Total cholesterol | CVDKP |
| rs3803578 | 2.82E-05 | Total cholesterol | CVDKP |
| rs889761 | 2.96E-05 | Total cholesterol | CVDKP |
| rs2926772 | 3.29E-05 | Total cholesterol | CVDKP |
| rs889765 | 3.77E-05 | Total cholesterol | CVDKP |
| rs113974433 | 5.46E-05 | Total cholesterol | CVDKP |
| rs71158757 | 5.53E-05 | Total cholesterol | CVDKP |
| rs2341001 | 7.18E-05 | Total cholesterol | CVDKP |
| rs3848236 | 7.61E-05 | Total cholesterol | CVDKP |

### SI Table S2

| Gene Name | Forward | Reverse |
|---|---|---|
| *Rps29* | GTCTGATCCGCAAATACGGG | AGCCTATGTCCTTCGCGTACT |
| *Abcb11* | CTGCCAAGGATGCTAATGCA | CGATGGCTACCCTTTGCTTCT |
| *Apob100* | AAGCACCTCCGAAAGTACGTG | CTCCAGCTCTACCTTACAGTTGA |
| *Asbt* | ACCACTTGCTCCACACTGCTT | CGTTCCTGAGTCAACCCACAT |
| *Abcg5* | TGCCCATTCCTTTAAAAATCC | GATGAACTGGACCCCTTGG |
| *Abcg8* | GGGGCTGATGCAGATTCA | GTAGCTGATGCCGATGACAA |
| *Acat2* | TCTTCTTCGCCTTCCTGCACTG | GAAGTCGAGTTCCACCAATCCC |
| *Cd36* | GAGCAACTGGTGGATGGTTT | GCAGAATCAAGGGAGAGCAC |
| *Cyp7a1* | CAGGGAGATGCTCTGTGTTCA | AGGCATACATCCCTTCCGTGA |
| *Dgat2* | GCGCTACTTCCGAGACTACTT | GGGCCTTATGCCAGGAAACT |
| *Fabp1* | ATGAACTTCTCCGGCAAGTACC | CTGACACCCCCTTGATGTCC |
| *Fasn* | TCCTGGGAGGAATGTAAACAGC | CACAAATTCATTCACTGCAGCC |
| *Fgf15* | ACGTCCTTGATGGCAATCG | GAGGACCAAAACGAACGAAATT |
| *Hmgcr* | AGCTTGCCCGAATTGTATGTG | TCTGTTGTGAACCATGTGACTTC |
| *Ldlr* | GCATCAGCTTGGACAAGGTGT | GGGAACAGCCACCATTGTTG |
| *Mttp* | ATACAAGCTCACGTACTCCACT | TCCACAGTAACACAACGTCCA |
| *Npc1l1* | CAACATCTTCATCTTTGTTCTTGAG | GCAATGTGAGCCTCTCG |
| *Nr1h3* | AGAGATGTCCTTGTGGCTGGAG | TCCACAACTCCGTTGCAGAATCAG |
| *Pcsk9* | GTGGTGATTGGATTGAGGCCATAG | CCCAAATGCATTGAGGGCCTTG |
| *Ppara* | TATTCGGCTGAAGCTGGTGTAC | CTGGCATTTGTTCCGGTTCT |
| *Ppard* | GACCAGAACACACGCTTCCT | CCGACATTCCATGTTGAGG |
| *Scarb1* | CGCCGACCCTGTGTTGTC | GGATGTCTAGGAACAAGGAATGCT |
| *Slc10a1* | ATGACCACCTGCTCCAGCTT | GCCTTTGTAGGGCACCTTGT |
| *Slc27a2* | ACAACATTCGTGCCAAGTCTCT | CTCCTCCACAGCTTCTTGTAGATC |
| *Vldlr* | TCCTGATTGCGAAGACGGTTCTG | ATGCGGCATGTTCTCATATGGC |
